# Nucleolar stress causes the entry into replicative senescence in budding yeast

**DOI:** 10.1101/297093

**Authors:** Sandrine Morlot, Song Jia, Isabelle Léger-Silvestre, Audrey Matifas, Olivier Gadal, Gilles Charvin

**Affiliations:** Developmental Biology and Stem Cells Department, Institut de Génétique et de Biologie Moléculaire et Cellulaire, Strasbourg, France; Centre National de la Recherche Scientifique, Illkirch, France; Institut National de la Santé et de la Recherche Médicale, Illkirch, France; Université de Strasbourg, Illkirch, France; Laboratoire de Biologie Moléculaire Eucaryote, Centre de Biologie Intégrative (CBI), Université de Toulouse, CNRS, UPS, Toulouse 31000, France

**Keywords:** Live cell imaging, microfluidics, ribosome biogenesis, replicative aging

## Abstract

The accumulation of <u>E</u>xtrachromosomal <u>r</u>DNA <u>C</u>ircles (ERCs) and their asymmetric segregation upon division have been hypothesized to be responsible for replicative senescence in mother yeasts and rejuvenation in daughter cells. However, it remains unclear by which molecular mechanisms ERCs would trigger the irreversible cell cycle slow-down leading to cell death. We show that ERCs accumulation is concomitant with a nucleolar stress, characterized by a massive accumulation of pre-rRNAs in the nucleolus, leading to a loss of nucleus-to-cytoplasm ratio, decreased growth rate and cell-cycle slow-down. This nucleolar stress, observed in old mothers, is not inherited by rejuvenated daughters. Unlike WT, in the long-lived mutant *fob1∆*, a majority of cells is devoid of nucleolar stress and does not experience replicative senescence before death. Our study provides a unique framework to order the successive steps that govern the transition to replicative senescence and highlights the causal role of nucleolar stress in cellular aging.

## Introduction

Budding yeast cells undergo a limited number of asymmetrical divisions before entering senescence and eventually dying, a phenomenon known as replicative aging (Mortimer and Johnston, 1959). Unlike symmetrically dividing unicellular organisms (Nakaoka and Wakamoto, 2017; Spivey et al., 2017), the replicative lifespan (RLS) in budding yeast follows a broad Gaussian distribution within a population, indicating that cell death is not a stochastic process, yet presents large cell-to-cell variability. Importantly, while the replicative age of mothers increases at each division, new born daughter cells recover a full replicative potential (Kennedy et al., 1994). Such rejuvenation of daughter cells led to postulate that cell death results from the accumulation of aging factors in mothers, which are asymmetrically segregated upon cell division (Egilmez and Jazwinski, 1989).

The accumulation of <u>E</u>xtrachromosomal <u>r</u>ibosomal DNA <u>C</u>ircles (ERCs) in aging mothers was the first potential aging factor that was studied in detail (Sinclair and Guarente, 1997). ERCs result from the excision of repeats from the rDNA cluster located on chromosome XII. This genomic region contains the highly transcribed 35S ribosomal RNA genes in tandem repeats (approximately 150 copies). A replication fork barrier (RFB), where the protein Fob1 binds, prevents the collision between the replication fork and the RNA polymerase I transcription of 35S rRNA gene (Brewer and Fangman, 1988; Kobayashi, 2003). The repetitive nature of the rDNA cluster and the stalling of the DNA replication complex at the RFB, which favors double strand breaks, promote recombination events. ERCs were shown to progressively accumulate in aging cells due to the presence of an <u>a</u>utonomously <u>r</u>eplicating <u>s</u>equence (ARS) on each rDNA repeat, that ensures the amplification of the excised DNA circles over successive divisions (Sinclair and Guarente, 1997). Strains deleted for the *FOB1* gene present lower amount of ERCs and an extended lifespan (Defossez et al., 1999) which further correlates accumulation of ERCs with replicative aging. Finally, to explain daughter rejuvenation with the ERC accumulation model, it was proposed that a diffusion barrier at the bud neck prevents the ERCs from being inherited by the daughter cells during cell division (Denoth-Lippuner et al., 2014; Shcheprova et al., 2008).

However, maintaining low ERC levels by genetically decreasing the recombination probability at the rDNA locus is not enough to promote longevity (Ganley et al., 2009). Furthermore, it is not clear why ERC accumulation is toxic to the cell and how it would lead to an arrest of proliferation (Ganley and Kobayashi, 2014). Another hypothesis favors rDNA instability, in particular the activity of the non-coding bidirectional promoter, called Epro, which is responsible for the amplification of rDNA repeats, rather than ERCs accumulation as being involved in replicative aging (Kobayashi, 2008; Saka et al., 2013). Hence, the mechanism that links rDNA regulation and replicative aging remains to be elucidated.

The dynamics of aging in budding yeast has been previously characterized by an abrupt transition, called the <u>S</u>enescence <u>E</u>ntry <u>P</u>oint (SEP), between an healthy state with regular and robust divisions and a senescent-like state where cell cycles are much longer and variable (Fehrmann et al., 2013). The molecular pathways responsible for this abrupt cellular transition are still not understood. In particular, it is not known how ERCs, rDNA instability and nucleolar activity, good candidates for aging factors, are linked to the SEP.

In this study, we analyzed in vivo the aging process in single yeast cells over multiple generations using microfluidic tools and time lapse microscopy. We show that a dramatic nucleolar stress combined with the accumulation of ERCs precede the SEP. This nucleolar stress is characterized by an upregulation of rDNA transcription and an accumulation of pre-rRNAs in the nucleolus followed, by an enlargement of the nucleus concomitantly with the irreversible cell cycle slow-down. Altogether, these findings highlight that the nucleolar steps of ribosome biogenesis play a critical role in the entry of replicative senescence.

## Results

### The number of rDNA repeats and the nucleolar volume increase before the irreversible cell cycle slow-down leading to cell death

To follow the real-time dynamics of the rDNA cluster during replicative aging in individual yeast cells, we used a previously developed strain (Miyazaki and Kobayashi, 2011), in which LacO sequences are inserted in all repeats of the rDNA array (Fig1A and Movie S1). In this strain, the protein Net1, involved in nucleolar silencing, telophase exit and stimulation of RNA Pol I-mediated transcription, is labelled with mCherry, which allows detecting the nucleolus. Thus, by monitoring GFP-LacI signal within the nucleolus delimited by NET1-mCherry fluorescence, we could estimate the evolution of rDNA repeats number in single mother cells trapped from birth to death in a microfluidic device (Fig1A), as previously described (Goulev et al., 2017). The <u>s</u>enescence <u>e</u>ntry <u>p</u>oint (SEP) is a particularly critical time point in yeast lifespan as it corresponds to the abrupt cell cycle slow-down occurring before cell death (Fig1B and 1C) (Fehrmann et al., 2013). We thus analyzed fluorescence signals after SEP alignment rather than birth alignment which blurs the cell cycle dynamics and does not allow to order cellular events (FigS1A). We observed that the total GFP-LacI fluorescence within the nucleolus increased significantly by 60% within the 5 divisions preceding SEP and further increased to 4.4-fold 5 divisions after SEP (Fig1D). If considering 150 repeats as the basal size of the rDNA cluster, fluorescence measurements allow us to estimate an increase to 240 repeats before SEP and to more than 600 repeats after SEP. This increase is likely to originate from ERCs accumulation as wild type cells hardly accumulate more than 200 repeats within the rDNA cluster (Ide et al., 2013). In agreement with this result, we also found an increased signal of FOB1-GFP before SEP (FigS1B), which suggests a higher rate of blocked replication forks at RFB before SEP and thus a higher rate of ERCs production.

**Figure 1:**
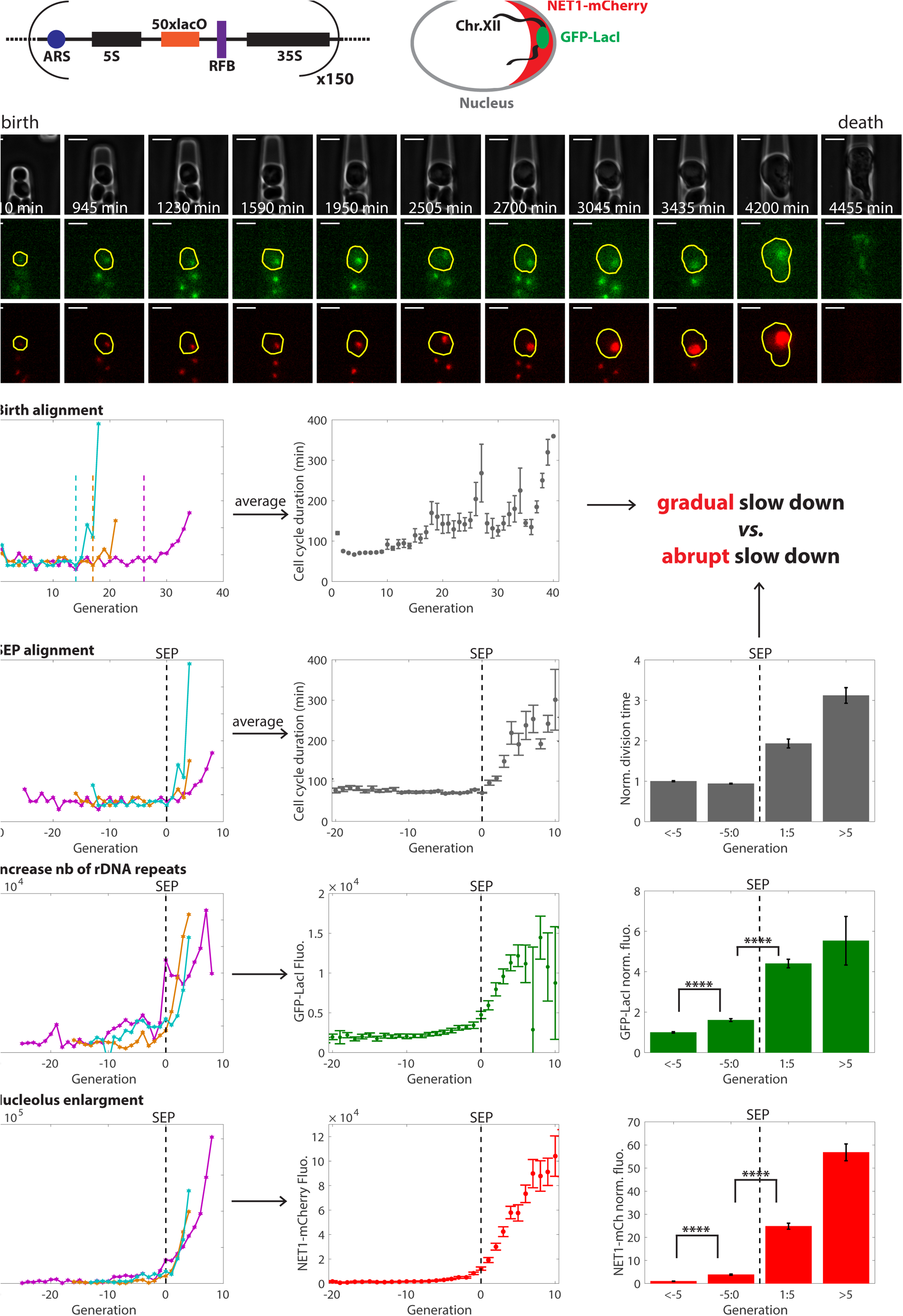
The number of rDNA repeats and the nucleolus size increase 5 divisions prior to SEP. (A) Top: sketch of LacO insertion on each rDNA repeat in a strain containing NET1-mCherry and GFP-LacI. Bottom: pictures of a mother cell, delimited by a yellow contour in the fluorescence channels, with GFP-LacI and NET1-mCherry markers, trapped in a cavity in a microfluidic device from birth to death. Scale bar: 5µm. (B) Left: duration of successive cell cycles for 3 single cells (magenta, cyan and orange). Middle: Average cell cycle duration as a function of age for single cell trajectories aligned from birth. (C) Left: same 3 single cells as in B aligned from SEP. Middle: Average cell cycle durations after SEP alignment. Right: normalized cell cycle duration for 4 classes of age: (1): more than 5 divisions before SEP, (2): 5 to 0 divisions before SEP, (3): 1 to 5 divisions after SEP and (4): more than 5 divisions after SEP; data are normalized to the average of the 1st class. (D) Total GFP-LacI fluorescence (arbitrary units) within the contour delimited by NET1-mCherry signal for the same 3 single cells as above (left) and averaged after SEP alignment (middle). Right: normalized GFP-LacI fluorescence for 4 classes of age. (E) Total NET1-mCherry fluorescence (arbitrary units) for the same 3 single cells as above (left) and averaged after SEP alignment (middle). Right: normalized NET1-mCherry fluorescence for 4 classes of age. N=49 mother cells (for all averaged curves).

Interestingly, NET1-mcherry signal increased to a larger extent than GFP-LacI before (3.9-fold) and after SEP (25-fold) (Fig1E and FigS1E). The nucleolar size has been shown to correlate positively with the cellular volume (Jorgensen et al., 2007). However, the sudden increase in NET1-mCherry signal was not due to an abrupt cell size enlargement as the cell area dynamics did not vary much during the SEP transition (FigS1C). This increase of NET1-mCherry fluorescence was mainly due to an enlargement of the nucleolar volume as NET1-mCherry mean fluorescence increases only by 36% after SEP (FigS1E) which was much lower than the 17-fold increase of the segmented area (FigS1D).

These results highlight that the nucleolus undergoes two major modifications prior to entry into cellular senescence: an increased copy number of ribosomal RNA genes and a volume expansion.

### A massive upregulation of rDNA transcription and of pre-rRNAs processing precedes the entry into senescence

We found a dramatic increase in the nucleolar volume occurring before the SEP. The nucleolus is the nuclear region were pre-rRNAs are synthetized and processed. Therefore, we investigated how rDNA transcription and pre-rRNAs levels evolve during the SEP transition.

To this end, we first monitored the total amount of RNA Pol1 by following its largest subunit RPA190, fused to GFP, throughout yeast lifespan (Fig2A and Movie S2). We measured an increase by 70% in RPA190-GFP total fluorescence over the 5 divisions preceding SEP, followed by a larger increase (up to 7-fold) after SEP (Fig2B). Next, to establish if this increased level of RNA Pol1 led to higher transcription at the rDNA locus, we measured the amount of pre-rRNAs in the nucleolus by performing RNA FISH on aging cells directly in the microfluidic chip (referred to as “FiSH on Chip” in the following). For this, we stopped the time-lapse experiment after 60 hours acquisition, when the microfluidic device was fully loaded and the ages of trapped mother cells were spread before and after SEP, as cells growing in cavities were not synchronized (left graphs Fig2C and 2D). We then performed RNA FISH staining on chip after cell fixation and cell wall digestion. The FISH probe targeted specifically the ITS1 region so that all pre-rRNAs from 35S to 20S would be labelled. FISH staining and RPA190-GFP signal presented a good colocalization, as expected (panels in Fig2C and 2D). For cells in a pre-SEP state (constant and short division timings), we measured a strong positive correlation (black dots in Fig2C, Pearson corr. coef. = 0.8) between the fluorescence levels of FISH probes and RPA190-GFP, indicating that the amount of pre-rRNAs scaled with the amount of RNA Pol1. For cells having already experienced SEP before fixation and FISH staining, the positive correlation remained, even though to a lower extent (magenta stars in Fig2C, Pearson corr. coef. = 0.5) (Fig2C), suggesting that the amount of RNA Pol1 and the rate of transcription could be uncoupled after entry into senescence. We measured a 4.4-fold increased amount of RNA Pol1 and a 2.6-fold increase of pre-rRNAs after the SEP transition (boxplot Fig2C). These results show that pre-rRNAs undergo a massive nucleolar accumulation upon the SEP.

**Figure 2:**
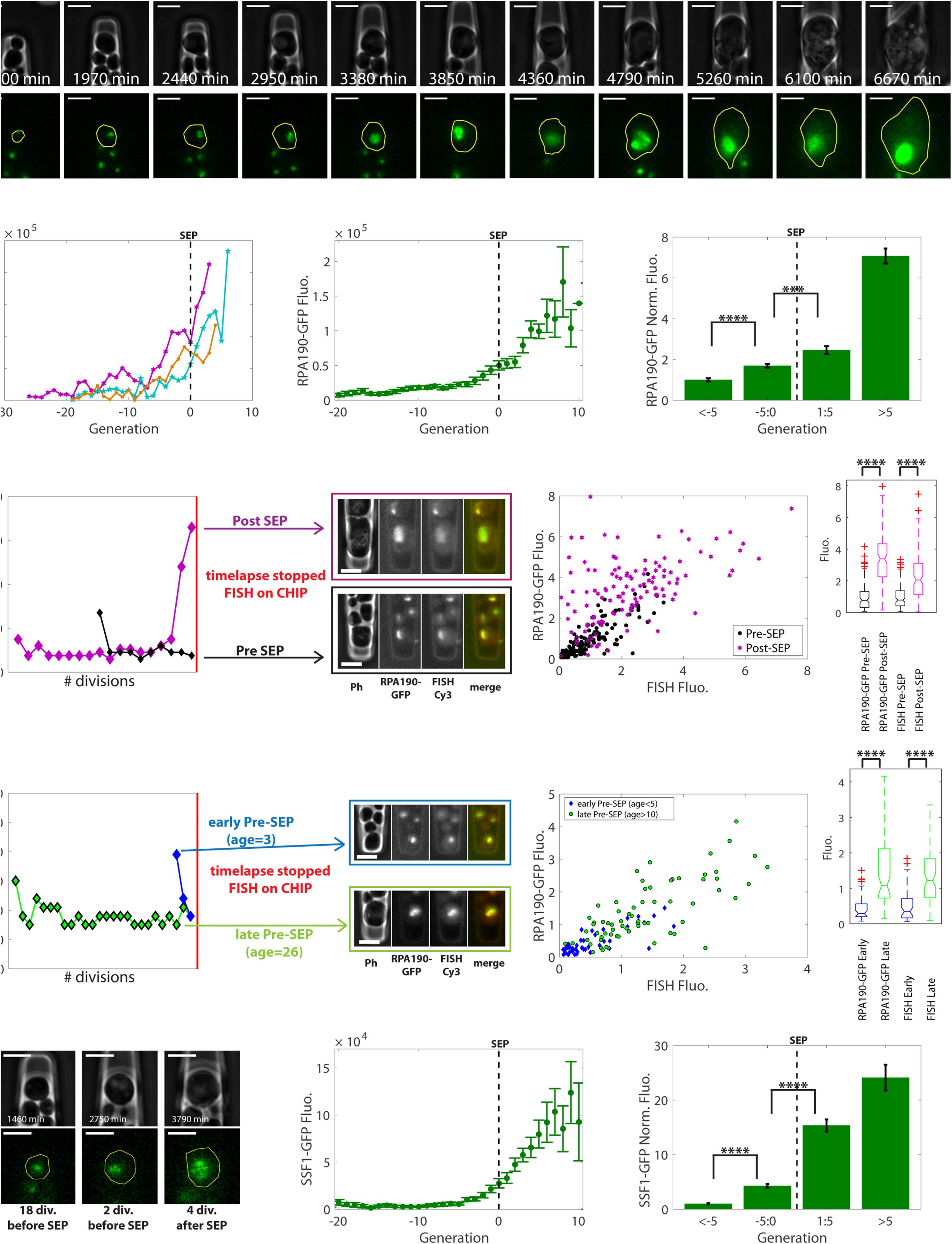
rRNAs synthesis and processing are upregulated before SEP. (A) Mother cell with RPA190-GFP marker in a microfluidic chip. (B) Total RPA190-GFP fluorescence (arbitrary units) for 3 single cells (left) and averaged after SEP alignment (middle). Right: normalized RPA190-GFP fluorescence for 4 classes of age. N=61 cells. (C) and (D) FISH on CHIP. Left: examples of cell cycle trajectories and corresponding pictures of cells labeled by RNA FISH directly on the microfluidic chip after 60 hours time-lapse acquisition. Middle: RPA190-GFP fluorescence as a function of FISH probe fluorescence. Right: boxplot of relative fluorescence of RPA-190 and FISH probe for pre-SEP cells (N= 136 cells, black in C), post-SEP cells (N= 123 cells, magenta in C), early pre-SEP cells (N= 45 cells, blue in D) and late pre-SEP cells (N= 68 cells, green in D). (E) Left: mother cell with SSF1-GFP marker in a microfluidic chip. Middle: total SSF1-GFP fluorescence averaged after SEP alignment. Right: normalized SSF1-GFP fluorescence for 4 classes of age. N= 56 cells. Scale bar: 5µm (for all pictures).

As RNA Pol1 showed an increasing amount before SEP (Fig. 2A), we tested whether pre-rRNAs levels followed a similar trend. To address this question, we focused on pre-SEP cells (black dots on Figure 2C) and compared ‘early pre-SEP’ cells (blue diamonds in Fig2D) having experienced less than 5 divisions with ‘late pre-SEP’ cells (green dots in Fig 2D) having undergone more than 10 divisions. ‘Late pre-SEP’ cells are more likely than ‘early pre-SEP’ cells to be close to the SEP and thus to present a higher level of RPA190-GFP. Indeed, we measured a 3.8-fold increase of RPA190-GFP fluorescence in the ‘late pre-SEP’ cells (boxplot Fig2D). We observed a good correlation between RNA Pol1 amount and FISH fluorescence in the 2 subgroups (Pearson corr. coeff. = 0.8 for ‘early pre-SEP’ and 0.7 for ‘late pre-SEP’, Figure 2D). Moreover, we measured a 3.9-fold increase in the pre-rRNAs levels in the ‘late pre-SEP’ cells, similar to the 3.8-fold increase in Pol1 levels (Fig2D). These results show that pre-rRNAs levels follow a dynamics very similar to RNA Pol1 before SEP. We thus conclude that the onset of accumulation of pre-rRNAs in the nucleolus precedes the entry into replicative senescence.

As the RNA FISH experiment could not discriminate between the successive transient pre-rRNAs from 35S to 20S, we could not determine whether rRNA processing was upregulated before SEP. Since SSF1 is a component of the 66S pre-ribosomal particles, we hypothesized that the level of this protein should follow the increase in pre-rRNA levels if rRNA processing was upregulated accordingly upon the SEP. Therefore, we monitored SSF1 protein fused to GFP throughout aging (Fig2E and Movie S3). Similarly to pre-rRNA levels, we observed that SSF1-GFP fluorescence increased by 4.2-fold before and up to 24-fold after SEP (Fig2E). These results show that both co- and post-transcriptional processing of pre-rRNAs are upregulated during the 5 divisions before SEP. These results suggest that the entry into senescence is preceded by a massive increase in the demand of ribosomal biogenesis that goes well beyond physiological growth requirements and results in the accumulation of pre-rRNAs in the nucleolus.

### The up-regulation of pre-rRNAs synthesis does not lead neither to an increase in ribosomes levels nor to a higher growth rate

Since senescent cells undergo a large accumulation of pre-ribosomes, we wondered whether this process was ultimately accompanied by a larger production of ribosomes and, eventually, an increase of the growth rate. First, we sought to evaluate if the nuclear export of pre-RNAs was following the same dynamics as the earlier steps of ribosome biogenesis.

To this end, we reasoned that the export rate of pre-rRNAs should scale with the amount of NOG2, which is a protein required for late pre60S ribosome maturation in the nucleoplasm and nuclear export. By monitoring the fusion NOG2-GFP (Fig3A and Movie S4), we measured a good scaling of fluorescence levels with the cell area before SEP (Fig3B and FigS2A), in sharp contrast with the levels of RPA190-GFP and SSF1-GFP, which increased much faster than cell area from 5 divisions before SEP (Fig3G, 3H, S2D and S2E). Hence, our results suggest that the constant basal rate of nucleoplasmic maturation of pre-ribosomes is uncoupled to the upregulation of the nucleolar steps of ribosome biogenesis.

**Figure 3:**
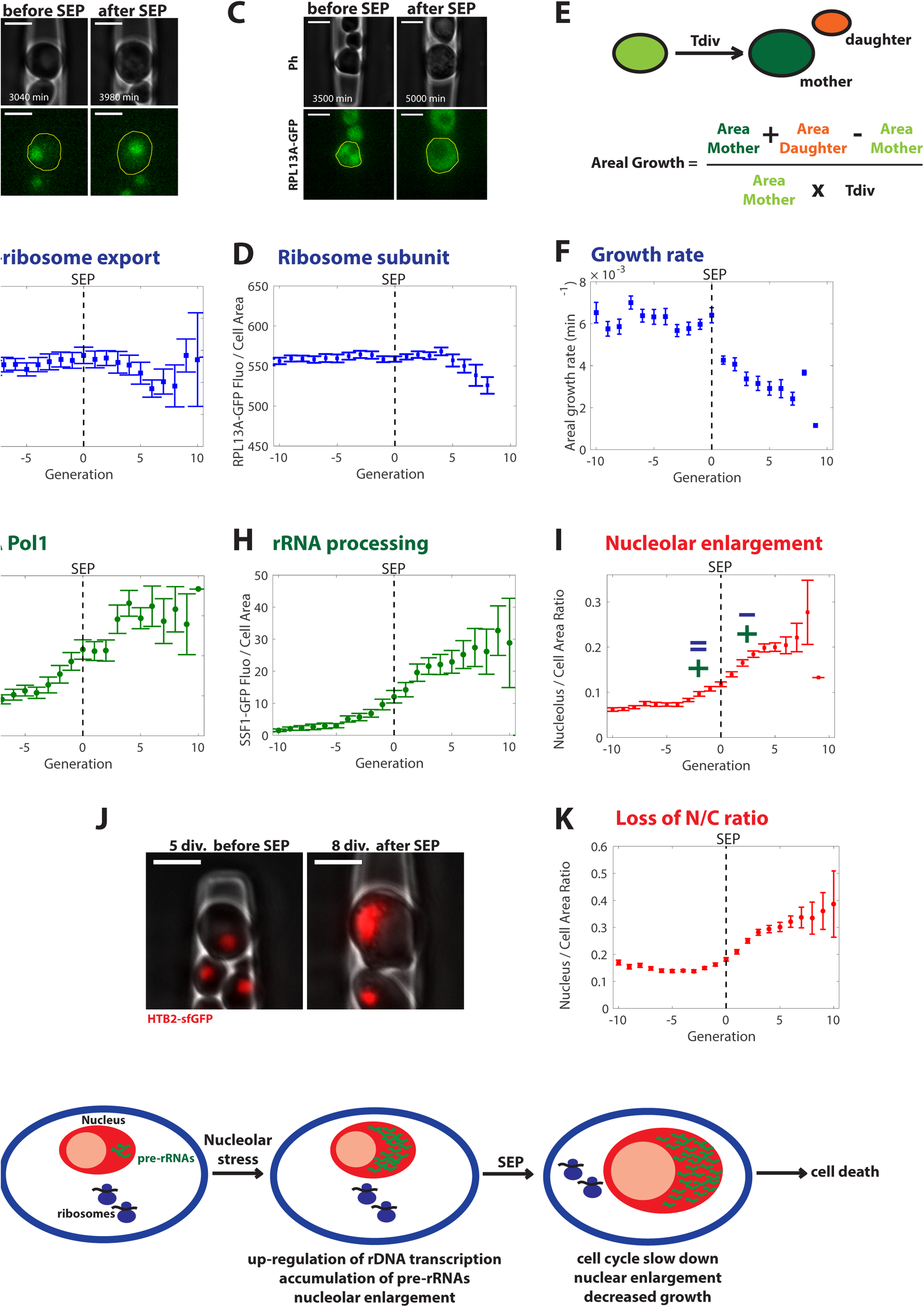
Ribosome concentration and growth rate do not increase during the entry into senescence. (A) Mother cell with NOG2-GFP marker on a microfluidic chip before and after SEP. (B) Ratio of NOG2-GFP total fluorescence over cellular area as a function of generation, averaged after SEP alignment. N= 33 cells. (C) Mother cell with RPL13A-GFP marker on a microfluidic chip before and after SEP. (D) Ratio of RPL13A-GFP total fluorescence over cellular area as a function of generation, averaged after SEP alignment. N= 32 cells. (E) Sketch explaining the calculation of growth rate based on segmented contour of the mother cells before budding (light green), the mother cell at the end of the division (dark green) and the new born daughter (orange). (F) Growth rate (measured from projected area) as a function of generation, averaged after SEP alignment. N=36 cells. (G) Ratio of RPA190-GFP total fluorescence over cellular area as a function of generation averaged after SEP alignment. N= 61 cells. (H) Ratio of SSF1-GFP total fluorescence over cellular area as a function of generation averaged after SEP alignment. N= 56 cells. (I) Ratio of nucleolar area (based on NET1-GFP segmentation) over cellular area as a function of generation, averaged after SEP alignment. N= 44 cells. (J) Mother cell with HTB2-sfGFP marker on a microfluidic chip before and after SEP. (K) Ratio of nuclear area (based on HTB2-sfGFP segmentation) over cellular area (or N/C ratio) as a function of generation, averaged after SEP alignment. N= 42 cells. (L) Model: nucleolar stress precedes SEP during replicative aging. Scale bar: 5µm (for all pictures).

Next, we investigated whether the cytoplasmic level of ribosomes was modified during the SEP transition by following RPL13A, a component of the 60S ribosomal subunit, fused to GFP (Fig3C and Movie S5)(Janssens and Veenhoff, 2016). Similarly to NOG2-GFP, we observed a constant ratio of RPL13A-GFP over the cellular area before SEP (Fig3D and FigS2B), indicating that the cytoplasmic concentration of ribosomes remained constant during the SEP transition, in striking contrast with the massive upregulation of early steps of ribosome biogenesis in the nucleolus.

Finally, we estimated the growth rate based on the segmented contours in phase contrast of mother and daughter cells (Fig3E). In healthy cells, the activation of ribosome biogenesis in the nucleolus normally responds to a higher demand in growth. In contrast, in aging cells experiencing a nucleolar expansion, we observed a constant growth rate until the SEP (Fig3F and FigS3C) in agreement with the constant concentration of ribosomes. Strikingly the growth rate started to significantly decrease at the SEP (Fig3F and FigS2C), while this dynamics is not noticeable for RPL13A-GFP and NOG2-GFP where a slight decrease occurred only several divisions after SEP (Fig3B and 3D). These results suggest that senescent cells undergo a nucleolar stress where nucleolar ribosome biogenesis, cytosolic ribosome levels and growth rate are uncoupled.

### The SEP is concomitant with a large increase in nucleoplasmic content

As the nucleolar volume drastically increased before SEP compared to the cell volume (Fig3I), we looked at the dynamics of the whole nucleus. We measured the nucleus size with the nuclear reporter HTB2-sfGFP (Fig3J and Movie S6). We observed an abrupt increase in nuclear content, concomitantly with the SEP (Fig4D). This increase resulted in a striking loss of the nuclear to cellular volume ratio or N/C ratio (Fig3K and 4E). Importantly, this behavior is not specific to histones, as we observed the same dynamics with the nuclear reporter NLS-sfGFP under the control of *ACT1* promoter (FigS2F). Using the NLS-sfGFP reporter, the increase in N/C ratio occurred slightly before the SEP. We explain this difference by the fact that, with the NLS-sfGFP reporter, we followed the whole nucleus including the nucleolus, whereas with HTB2-sfGFP, we mainly measured the nucleoplasmic volume, as the nucleolus contains a low level of histones. Removing the NLS sequence abolished the sudden increase in sfGFP fluorescence at SEP (FigS2G), further highlighting the specific dynamics of the nucleus. The increase in N/C ratio is also peculiar when compared to the other organelles such as the vacuoles. Indeed using the vacuolar marker VPH1-GFP, we measured that the vacuole occupies a constant fraction of 30% of the cellular area throughout lifespan (FigS2H), while the nuclear area represents 10% of the cellular area before SEP and suddenly increases up to 40% after SEP (Fig3K, S2F, 4E). In addition, high N/C ratio is an unexpected feature of aging cells as this ratio is known to be strongly robust in wild type young yeast cells and in numerous mutants including cell size and cell cycle mutants (Jorgensen et al., 2007; Neumann and Nurse, 2007). Therefore the loss of N/C ratio, as a non-physiological characteristics, could be mechanistically linked to the irreversible cell cycle slow-down leading to cell death.

### Nucleolar stress is not transmitted to rejuvenated daughter cells

We have identified a nucleolar stress occurring in aging mother cells characterized by an upregulation, in the nucleolar steps, of ribosome biogenesis which is inefficient in producing more ribosomes and in increasing growth (Fig3L). This event was found to be tightly related to – and even preceded- the cell cycle slow-down that defines the SEP. According to the yeast replicative aging paradigm, daughters of aging mothers recover a full replicative potential, as putative aging markers are not transmitted to daughter cells (Kennedy et al., 1994). In the following, we sought to investigate whether our model of entry into senescence triggered by a nucleolar stress was compatible with the previously reported daughter rejuvenation.

To this end, first, as a proxy for assessing the recovery of physiological function in new born daughters, we measured the division time of daughters of aging mothers. For this, we took advantage of the cavities in our microfluidic device which allowed tracking of the daughter lineage of trapped mother cells for a couple of divisions. Strikingly, we found that the daughter cells of post-SEP mothers recovered a cell cycle duration that was identical to pre-SEP cells (Fig4A). This revealed that one single division was sufficient to remove the deleterious effects of replicative aging on cell cycle progression.

**Figure 4:**
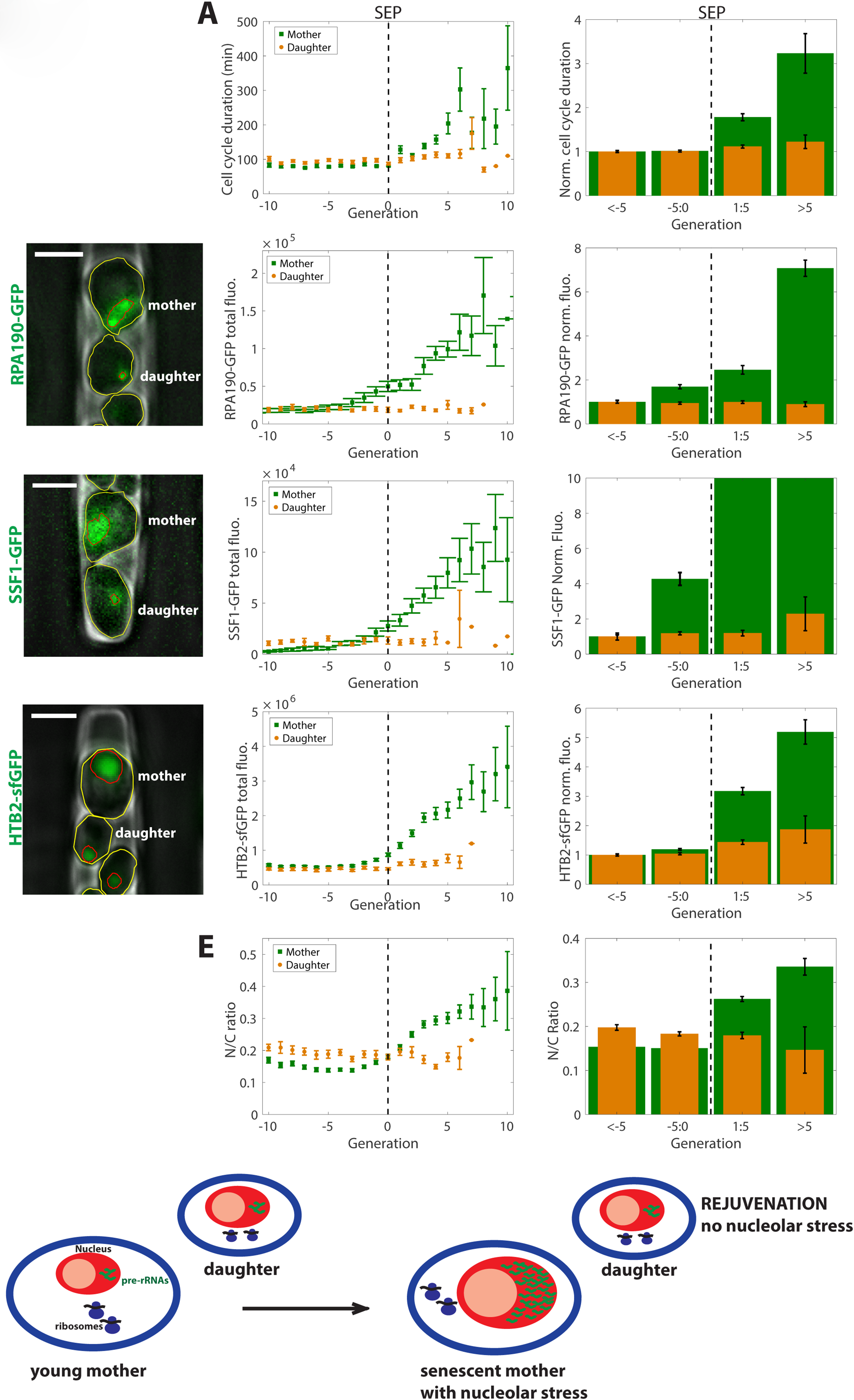
rRNA synthesis and nuclear size are rejuvenated to basal levels in daughter cells of aging mother. (A) Left: Cell cycle duration of mother (green) and daughter (orange) cells as a function of generation, averaged after SEP alignment. Right: normalized cell cycle duration of daughters (orange) and mothers (green) for 4 classes of mother age. N= 29 cells. (B) Left: picture of a senescent mother cell and its daughter cell with RPA190-GFP. Cell is delimited by a yellow contour and fluorescence signal by a red contour. Middle: RPA190-GFP total fluorescence in mothers (green) and daughters (orange) as a function of generation, averaged after SEP alignment. Right: RPA190-GFP normalized fluorescence in mother (green) and daughter (orange) cells for 4 classes of mother age. N= 21-61 cells. (C) Same as (B) for SSF1-GFP marker. N= 17-56 cells. (D) Same as (B) for HTB2-sfGFP marker. N=42 cells. (E) Left: N/C ratio of mother (green) and daughter (orange) cells as a function of generation, averaged after SEP alignment. Right: N/C ratio of daughters (orange) and mothers (green) for 4 classes of mother age. N= 42 cells. (F) Model: daughter cells of senescent mother do not inherit nucleolar stress. Scale bar: 5µm (for all pictures).

Then, we asked whether this rejuvenation process was driven by an asymmetrical segregation of the nucleolar stress markers upon division. Specifically, as RNA Pol1 and pre-rRNAs were both shown to increase in mother cells prior to SEP, we measured the fluorescence of RPA190-GFP and SSF1-GFP markers in the successive daughters of aging mother cells. Remarkably, daughter cells were born with a constant basal amount of these proteins even when the levels increased in mothers after the SEP (Fig4B and 4C). This demonstrated that daughter cells recovered basal levels of rDNA transcription and pre-rRNAs processing even when the mother acquired a strong senescence-associated impairment in nucleolar activity.

To further investigate the mechanism at stake in the segregation of age between mothers and daughters, we measured the nuclear size in daughters of aging mothers based on HTB2-sfGFP fluorescence. Importantly, we observed that daughters of post-SEP mothers inherited a nucleus of basal size despite the large increase in nuclear size in their mothers (Fig4D). Similarly, the N/C ratio was also rescued in daughter cells (Fig4E), further illustrating that daughters had recovered a normal physiology (Fig4F). Altogether, these results revealed the existence of an efficient rejuvenation mechanism based on asymmetrical partitioning of the nucleus/nucleolus upon division, which prevents the inheritance of nucleolar stress in new born daughters.

### Fob1 ties the onset of nucleolar stress to the entry into senescence

The fact that a nucleolar stress consistently occurs ahead of the onset of cell cycle slow-down upon entry into senescence argues in favor of a mechanistic link between these two events. To further establish their causality, we used the *fob1* mutant to perturb the stability of the rDNA and assess its consequence on the dynamics of entry into senescence.

The mutant fob1Δ has been shown to present a lifespan extension of 30% compared to wild type (Defossez et al., 1999). Measurements in our microfluidic device confirmed the enhanced longevity of this mutant (Table 1, Fig5A). Interestingly, a large majority (68.3%) of *fob1*Δ cells does not present any cell cycle slow-down before death while only 7% of WT cells have no SEP transition (Table1, Fig5B and 5C). These *fob1*Δ cells without SEP (referred to as ‘fob1Δ NO SEP’ in the following) present a further increased lifespan (RLS=37, Fig5A) than the fob1Δ cells experiencing a SEP (referred to as ‘*fob1*Δ with SEP’, RLS=34, still longer-lived than WT). Interestingly, in ‘*fob1*Δ with SEP’, the onset of cell cycle slow-down is delayed (Table1) but the dynamics of the cell cycle slow-down is similar to WT (Fig5D). It is worth noticing that WT, ‘fob1Δ with SEP’ and ‘fob1Δ NO SEP’ populations have similar cell area dynamics (FigS3A) when aligned from birth. When aligned from SEP, fob1Δ cells present a slightly larger area at SEP than WT cells (FigS3B) showing that SEP is not triggered by a cellular volume threshold. These results suggest that the observed differences with respect to the entry into senescence cannot be attributed to cell size effects and reinforce the idea that cell size is not an aging factor.

**Figure 5:**
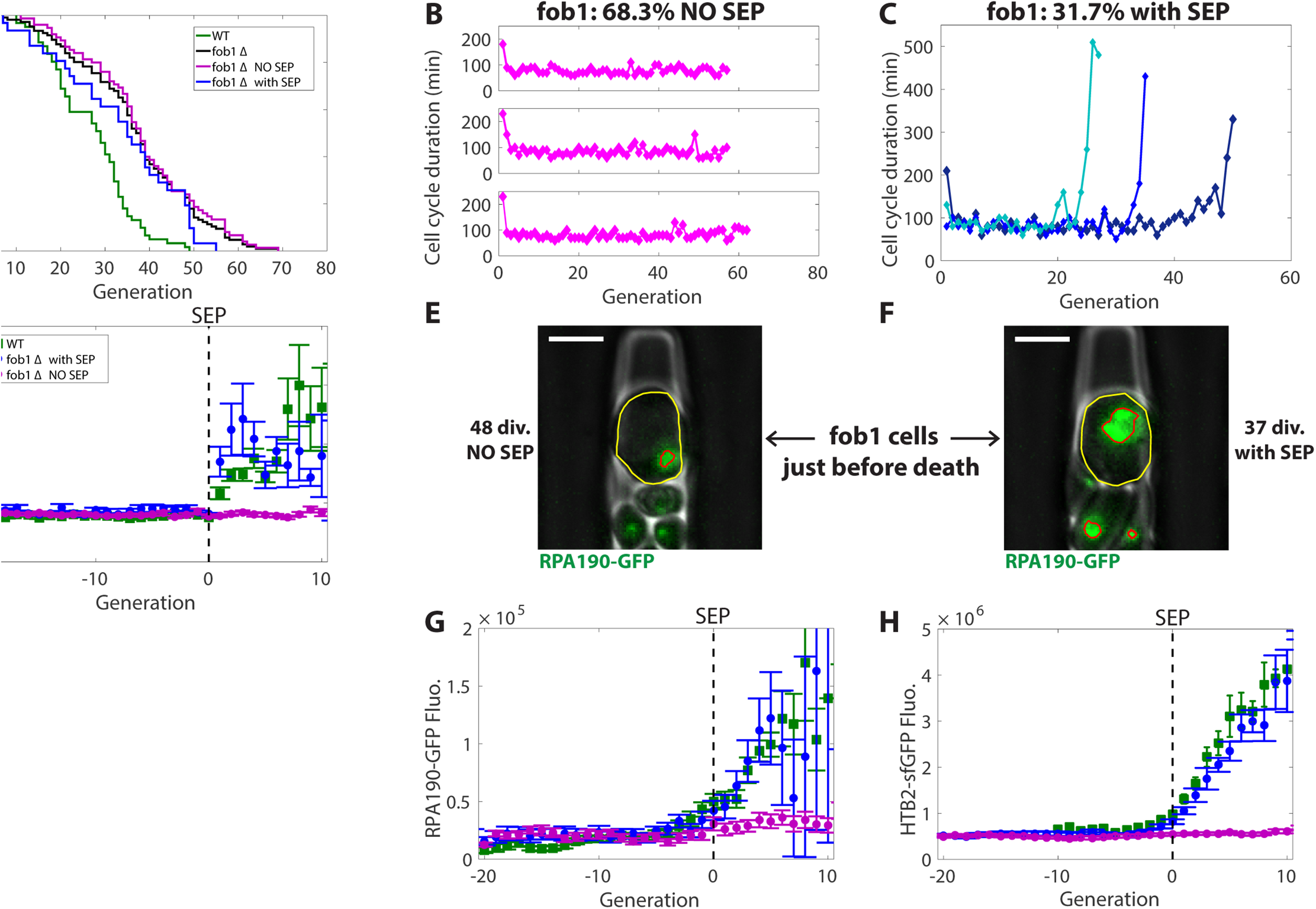
Senescence entry point results from a stochastic process dysregulating nucleolar activity. (A) Survival curves of fob1Δ strain (N= 126) and WT (N= 61). (B) and (C) Cell cycle durations for 3 single cells of fob1Δ strain without any SEP before death (B) and with SEP (C). (D) Cell cycle duration averaged after SEP alignment for WT (green, N= 57) and fob1Δ cells with SEP (blue, N= 40). Fob1Δ cells without SEP (magenta, N= 84) were aligned from birth and generation 0 was set to 30 (median SEP for fob1Δ cells with SEP). (E) and (F): Pictures of fob1Δ cells with RPA190-GFP marker, just before death, without a SEP (E) and with a SEP (F). Cell is delimited by a yellow contour and fluorescence signal by a red contour. Scale bar: 5µm. (G) RPA190-GFP fluorescence and (H) HTB2-sfGFP fluorescence in WT (green), “fob1 with SEP” (blue), “fob1 NO SEP” (magenta) after SEP alignment. N= 20-61 cells.

**Table 1:** aging parameters of the strains used in this study

| strain | N | RLS | % NO SEP | RLS NO SEP | RLS with SEP | SEP |
| --- | --- | --- | --- | --- | --- | --- |
| BY4742 | 61 | 29 | 7 | 35 | 28 | 20 |
| fob1Delta | 126 | 36 | 68.3 | 37 | 34 | 30 |
| NET1-GFP | 51 | 19 | 9.8 | 14 | 19 | 14.5 |
| fob1D+NET1-GFP | 135 | 43 | 58 | 48 | 37 | 30 |
| RPA190-GFP | 78 | 19 | 11.5 | 13 | 19 | 14 |
| fob1D+RPA190-GFP | 101 | 34 | 64.4 | 39 | 22.5 | 21 |
| HTB2-sfGFP | 112 | 27.5 | 18.75 | 39 | 26 | 19 |
| fob1D+HTB2-sfGFP | 56 | 39 | 62.5 | 45 | 33 | 27 |

Altogether, these results indicate that a common mechanism may be at stake in the progression through senescence *(i.e*. in the post-SEP period of the lifespan) in both WT and *fob1*Δ mutant. However, it suggests that the SEP trigger is a stochastic event, the probability of which is determined by Fob1, presumably through the control of the rDNA instability.

To further check this hypothesis, we analyzed the fluorescence of RPA190-GFP, subunit of RNA Pol1 in the two *fob1*∆ subpopulations. The population ‘*fob1*Δ with SEP’ experienced a significant increase of RPA190-GFP fluorescence before SEP similar to WT and cells died with a large amount of RNA Pol1 (Fig5F and Movie S8). In contrast, the ‘*fob1*Δ NO SEP’ population kept a basal level of RPA190-GFP throughout lifespan (Fig5G) and cells died with a physiological level of RNA Pol1 even after more than 30 divisions (Fig5E and Movie S7). Daughter cells of both populations ‘*fob1*Δ NO SEP’ and ‘*fob1*Δ with SEP’ recover similar levels of RPA190-GFP (FigS3C), showing that the maintenance of small nucleoli in ‘*fob1*Δ NO SEP’ is not due to an impaired asymmetrical nuclear segregation upon division but rather to a better homeostasis of nucleolar activity during aging.

Similarly, we measured the fluorescence of the nuclear marker HTB2-sfGFP in the two populations ‘fob1Δ NO SEP’ and ‘fob1Δ with SEP’. In agreement with the nucleolar markers, only the cells experiencing a SEP presented an increased level of HTB2-sfGFP very similar to WT (Fig5H). We also calculated the growth rate of *fob1*∆ cells. In agreement with the 2 previous senescence markers (cell cycle slow-down and increased nuclear size), ‘fob1Δ with SEP’ population presented a decreased growth rate at SEP similar to WT while ‘*fob1*Δ NO SEP’ cells maintained a constant growth rate throughout lifespan (FigS3D).

In conclusion, we show that the FOB1-dependent onset of nucleolar stress predicts, in the following 5 divisions, the entry into replicative senescence, characterized by sudden cell cycle slow down, decreased growth rate and nuclear enlargement.

## Discussion

By combining microfluidics, time-lapse microscopy and quantitative single cell analysis, we have uncovered that nucleolar stress is a predictive marker of imminent entry into senescence. Alignment from SEP was a key methodological breakthrough to chronologically order the molecular events leading to cell death. Indeed, when data were aligned from cell birth, as it is usually performed, all parameters gave similar gradual dynamics, which does not allow to distinguish causes from consequences (FigS1A). By aligning single cell trajectories from SEP, we identified that the nucleolar initiation of ribosome biogenesis is upregulated in a FOB1-dependent-way and is followed, within 5 divisions, by a massive nuclear enlargement, a decreased growth rate and the irreversible cell cycle slow-down, leading eventually to cell death (Fig3L and 4F). Thus, our results contradicts the classical paradigm of a progressive decline in physiological functions during aging and highlights the nucleolar stress as a critical step during replicative aging.

Ribosome biogenesis has been previously identified as a key pathway involved in yeast replicative aging. Indeed, in a genome-wide analysis of aged-sorted cell populations combining both mRNA sequencing and mass spectrometry, the authors showed that the transcription of protein biogenesis-related genes gets uncoupled, with age, from the translation of corresponding mRNAs (Janssens et al., 2015). They further suggested, based on computational analysis, that this uncoupling could likely cause replicative aging. In agreement with this study, we described a detailed molecular mechanism connecting the dysregulation of ribosome biogenesis, characterized by the uncoupling between pre-rRNAs synthesis and ribosomes maturation, to cellular aging. We established that a stochastic event triggers this nucleolar stress (and subsequently cellular senescence) almost systematically in WT cells and in a small fraction of *fob1*Δ cells. ERCs excision and/or rDNA cluster amplification are very likely to be this stochastic event as FOB1 regulates these processes (Defossez et al., 1999; Kobayashi et al., 1998). In wild type cells, ERCs are generated by the FOB1-dependent stalling of the replication fork at RFB, whereas in *fob1*Δ cells, ERCs are produced by the collision of the replication fork against RNA Pol1 transcription machinery (Takeuchi et al., 2003). This latter event occurs less frequently than FOB1-dependent excision as only half of 35S genes are actively transcribed (Dammann et al., 1993) and less than one third of ARS, in the rDNA cluster, are fired during S phase (Brewer and Fangman, 1988). Thus, some *fob1*Δ cells would never experience any ERC excision event during their lifespan. This would explain the two sub-populations “*fob1*Δ NO SEP” and “fob1Δ with SEP” that we identified. Our results also identified an atypical activation of rDNA transcription leading to cellular senescence. Indeed we show that the increased number of ribosomal RNA genes and the enhanced activity of RNA Pol1 are a unique feature of aging cells close to the senescence transition; as in healthy young yeast cells, RNA Pol1 activity is up-regulated, in a compensatory mechanism, only when the number of repeats in the rDNA cluster is reduced (Takeuchi et al., 2003).

We meticulously characterized the entry into senescence as the simultaneous onset of three sudden major cellular alterations: cell cycle slow down, decreased growth rate and nuclear enlargement. It still remains to be determined the precise interplay between these three physiological modifications and how they lead to cell death. The massive nuclear enlargement and the subsequent loss of N/C ratio at SEP are striking aging markers as the nuclear volume should normally robustly scale with the cellular volume (Neumann and Nurse, 2007). It has recently been shown that disrupting mRNA export machinery in *S*. *pombe* increases significantly the N/C ratio by accumulation of RNAs (Kume et al., 2017). In addition, premature rRNAs are known to be trapped within the nucleolus (Gadal et al., 2002). The transition from the nucleolus to the nucleoplasm is accompanied by major compositional changes in preribosome (Kressler et al., 2017). Thus, the dramatic accumulation of pre-rRNAs, trapped in the nucleolus, could probably result in the accumulation of further nuclear intermediates involved in ribosome biogenesis and be directly responsible for the sudden loss of N/C ratio during replicative aging. This abrupt increase in nuclear volume, together with the accumulation of nuclear proteins, might then trigger a pathway slowing down cell cycle to allow the cell to reach a volume matching its nuclear size. However, pre-ribosomal particles synthetized in senescent cells are not export-competent and remain in the nucleolus. Therefore, growth rate does not increase, thus cellular volume never catches up the appropriate physiological N/C ratio, reinforcing the activation of the pathway slowing down cell cycle, which could explain the irreversibility of the senescent state.

The nucleolar mechanism described in this study is likely conserved across species as similar observations were also reported in several studies in metazoans. In the germline stem cells of Drosophila male, the rDNA array on chromosome X is silenced in young flies but active in old flies (Lu et al., 2018). In mouse embryonic fibroblasts, oncogenic stress induces rRNA transcription and triggers cellular senescence (Nishimura et al., 2015). Our data precise that the transcription at the rDNA cluster is activated before the onset of cellular senescence in physiological conditions (no external induction of senescence). In particular, our work clearly demonstrates that nucleolar dysregulation is not a mere consequence of the aging process but appears before the establishment of a senescent state. More recently, it has been found that small nucleoli in post-mitotic cells are a hallmark of an extended longevity in *C. elegans*, Drosophila, Mice and human muscle tissues (Tiku et al., 2016). These results, together with our study, highlight the crucial role of the nucleolus in both replicative and chronological aging, suggesting the existence of a global and conserved nucleolar mechanism controlling longevity.

## Material and Methods

### Yeast strains, plasmids and media

All strains used in this study are congenic to S288C, except TMY8-BY4B (strain TMY8 back-crossed 4 times with BY4742/41). All GFP-labeled strains were provided from M. Knop GFP collection. Fob1Δ strain was purchased from Euroscarf, crossed with strains containing relevant GFP markers and genotyped by PCR. The HTB2-sfGFP, Act1pr-sfGFP and Act1pr-NLS-GFP strains were generated using DNA editing and yeast genetics techniques.

Prior to loading into microfluidic chips, freshly thawed cells were grown overnight and then diluted in the morning to allow a few divisions in exponential growth.

RNA FISH buffers:
-Buffer B: 54.66g sorbitol + 25ml Phosphate Buffer + 195ml sterile water (250ml final volume)
-Phosphate buffer: 136.09g KH2PO4 +228.23g K2HPO4 (1L final volume)
-Zymolase mix: 2ml buffer B + 4μL PMSF (stock: 0.1M) + 20μL Vanadium (stock: 200mM) + 4μL βME (stock: 14.3M pur) + 20μL Zymolase 100T (stock 5mg/ml in water)
-20x SSC: 175.3g NaCl + 82.3g Sodium tri Citrate + adjust to 1L with H2O
-Hybridation mix solution A: 4μl FISH probe (stock 10ng/μl) + 8μl pur formamide + 4μl 2xSSC + 4μl tRNA (stock 10mg/ml) + 14μl H2O
-Hybridation mix solution B: 2μl BSA (5% in 4xSSC) + 4μl Vanadium (stock: 200mM) + 40μL 4xSSC.

To prepare hybridation mix, solution A was incubated 5min at 98^°^C then added to solution B at RT.

FISH probe sequence: GCACAGAAATCTCTCACCGTTTGGAATAGCAAGAAAGAAACTTACAAGC with Cy3 dye in 5’. The probe targets ITS1 region between the A2 cleavage site and the 18S coding region.

### Microfluidics

The microfluidic master mold was made using standard soft-lithography techniques in the FEMTO-ST nanotechnology platform of the French Renatech network (Besançon, France). Prototypic molds were replicated in epoxy to ensure long-term preservation. The micro-channels were cast by curing PDMS (Sylgard 184, 10:1 mixing ratio) and then covalently bound to a 24 × 50 mm coverslip using plasma surface activation (Diener, Germany). The assembled chip was then baked 1 hour at 60°C to consolidate covalent bonds between glass and PDMS and then perfused with media circulating in Tygon tubing with a peristaltic pump (Ismatec, Switzerland) at a 10μL/min flow rate. After 2 hours of PDMS rehydration, yeast cells were loaded to the chip with a 1ml syringe and a 23G needle.

### Time-Lapse Microscopy

All experiments have been replicated at least twice.

Cells were imaged using an inverted Nikon Ti-E microscope. Fluorescence illumination was achieved using LED light (Lumencor) and light was collected using a 60× N.A. 1.4 objective and a CMOS camera Hamamatsu Orca Flash 4.0. We used an automated stage in order to follow up to 60 positions in parallel over the course of the experiment. Images were acquired every 10 or 15 min for a total duration of 140 hours (full lifespan) or 60 hours (RNA FISH) using NIS software. Focus was maintained with the Nikon Perfect Focus System. A constant temperature of 30°C was maintained on the chip using custom sample holder with thermoelectric modules, an objective heater with heating resistors and a PID controller (5C7-195, Oven Industries).

### Image Analysis

After acquisition, NIS raw data were analyzed using custom matlab software: phylocell and autotrack available on //github.com/gcharvin. Cell contours and fluorescent markers were segmented using a watershed algorithm and tracking was achieved with the Hungarian method.

### RNA FISH on CHIP

Time-lapse was stopped after 60 hours acquisition. 4% paraformaldehyde was perfused in the chip for 30 min at RT for cell fixation then the chip was washed with buffer B for 20min. Cell wall was digested by flowing zymolase mix for 20 min at RT followed by a 20min wash of buffer B. We then rinsed the chip with cold 70% ethanol for 5min, then 15min with 2xSSC, then 20min with 10% formamide in 2xSSC. Hybridation mix with 1ng/µl FISH probe was injected into the chip which was then kept at 37°C for 3hours. After hybridation, we rinsed the chip 30 min with warm (37^°^C) 10% formamide in 2xSSC, then 20 min with Triton X-100 (1% in 2xSSC) and finally 30min with 1xSSC. Cells in the chip were then imaged on a Nikon Ti-Eclipse with a mCherry filter to acquire FISH probe signal (50% led power, 300ms exposure time, binning 2x2) with a GFP filter for RPA190-GFP/NET1-GFP signal (20% led power, 100ms exposure time, binning 2x2) and in phase contrast.

## Acknowledgements

This work was partly supported by the French RENATECH network. We thank Denis Fumagalli and the MEDIAPREP facility of IGBMC for preparing media. We thank Prof. Takehiko Kobayashi for providing the strain TMY8. We are grateful to Theo Aspert, Basile Jacquel and Sophie Quintin for careful reading of the manuscript.

## Author contributions

SM, OG and GC designed the project. SM and SJ carried out research and analyzed data. SM and ILS developed the “FISH on chip” methodology. AM designed the yeast strains used in this study. SM wrote the manuscript.

---

**Figure S1, related to Figure 1.**
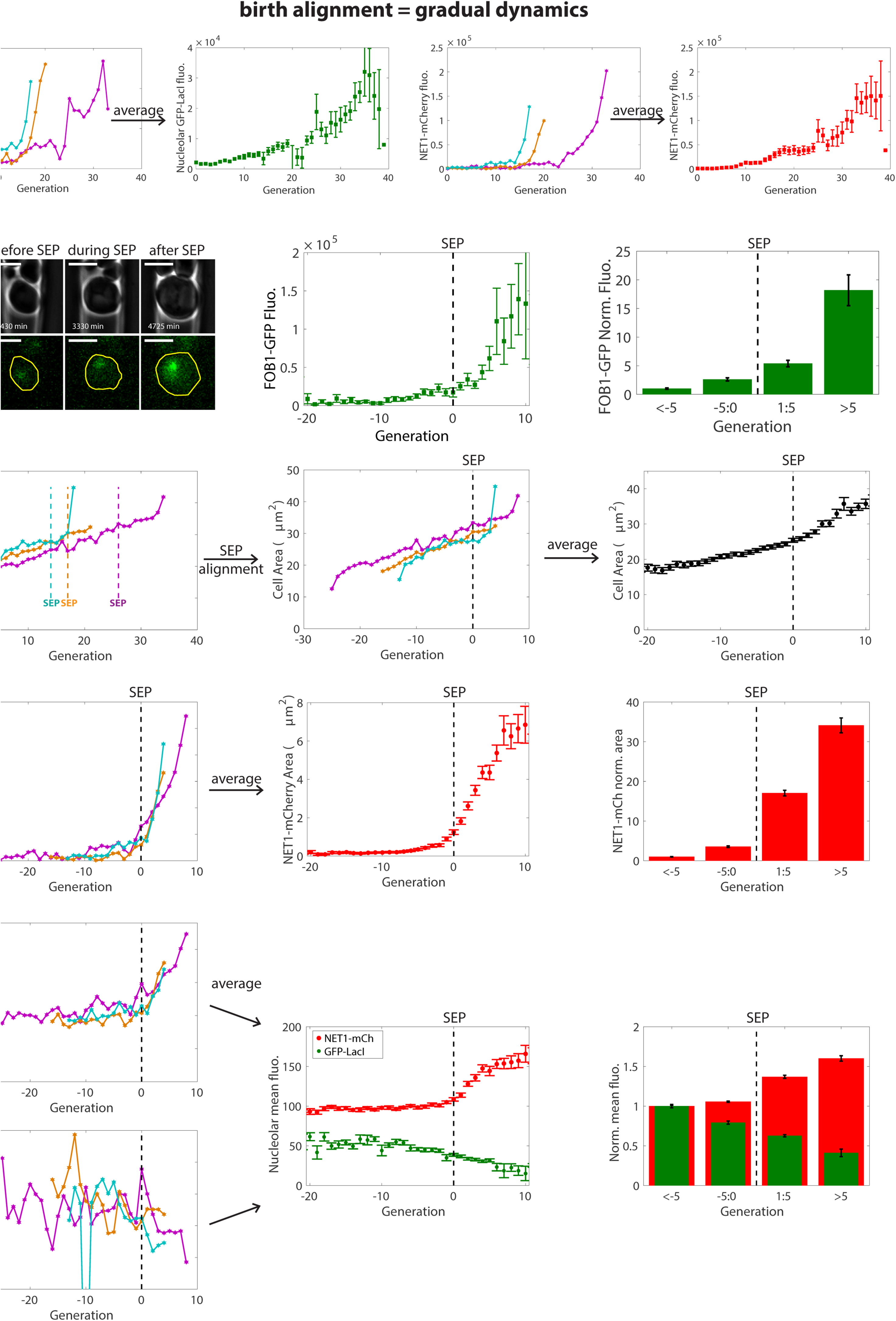
(A) From left to right: Total GFP-LacI fluorescence (arbitrary units) within the contour delimited by NET1-mCherry signal for the same 3 single cells as in Fig1, aligned from birth. Total GFP-LacI fluorescence averaged after birth alignment (N=49). Total NET1-mCherry fluorescence for the same 3 single cells as in Fig1, aligned from birth. Total NET1-mCherry fluorescence averaged after birth alignment (N=49). (B) Left: pictures of a mother cell with FOB1-GFP marker in a microfluidic chip. Middle: total FOB1-GFP fluorescence averaged after SEP alignment. Right: normalized FOB1-GFP fluorescence for 4 classes of age. N= 28 cells. Scale bar: 5µm. (C) Left: Cell area as a function of generation for 3 single cells (magenta, cyan, orange, same cells as in main Fig1) aligned from birth. Middle: Cell area for the same 3 single cells aligned from SEP. Right: Cell area as a function of generation, averaged after SEP alignment. (D) Left: segmented area of NET-mCherry signal for the same 3 single cells as above and main Fig1 aligned from SEP. Middle: averaged segmented area of NET1-mCherry after SEP alignment. Right: normalized NET1-mCherry segmented area for 4 classes of age. N= 49. (E) Left: NET1-mCherry (top) and GFP-LacI (bottom) mean nucleolar fluorescence for the same 3 single cells as a function of generation, after SEP alignment. Middle: averaged mean fluorescence of NET1-mCherry (red) and GFP-LacI (green) as a function of generation, after SEP alignment. Right: normalized mean fluorescence of NET1-mCherry (red) and GFP-LacI (green) for 4 classes of age. N= 49 cells.

**Figure S2, related to Figure 3.**
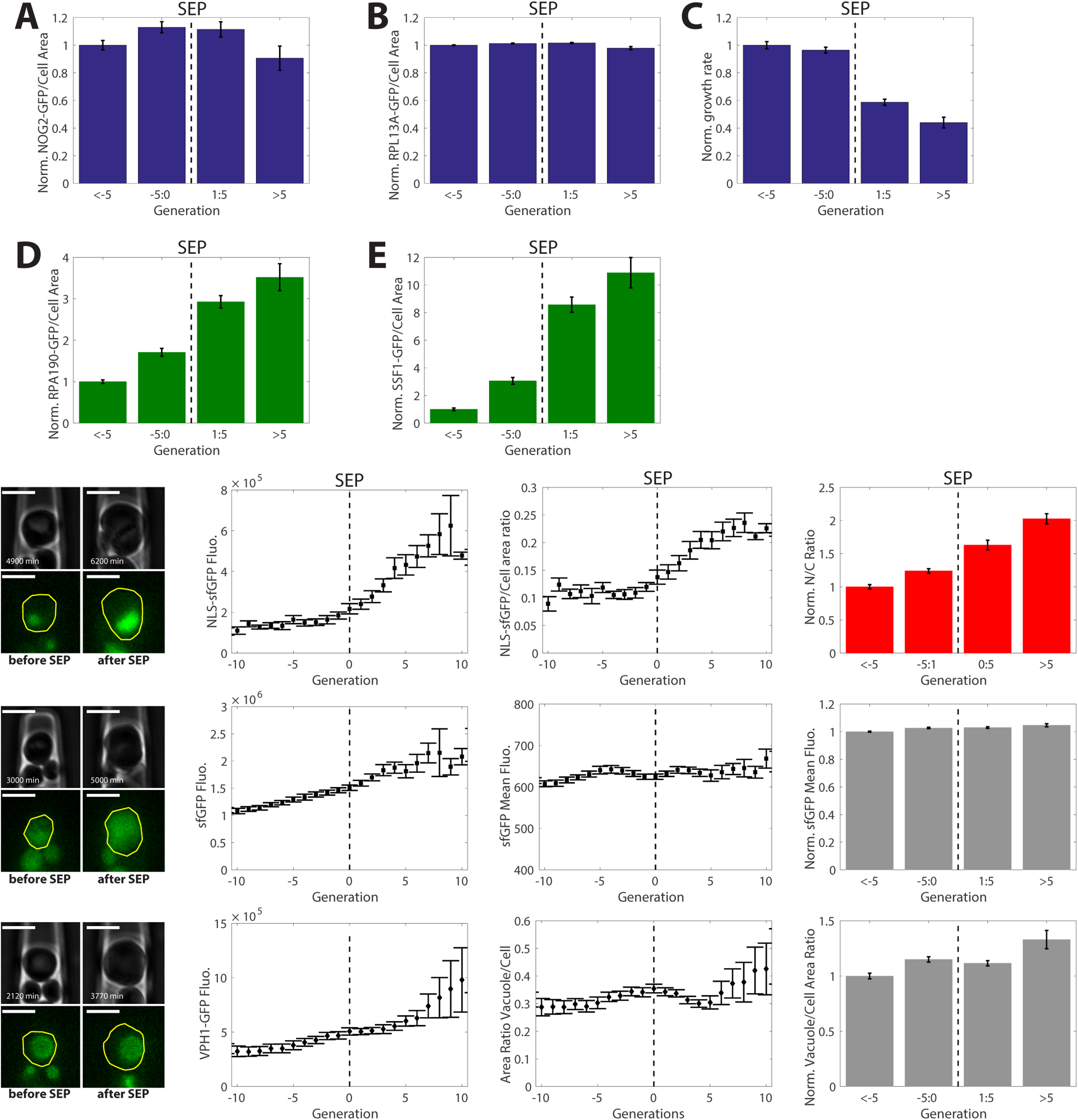
(A) Normalized ratio of NOG2-GFP total fluorescence over cellular area for 4 classes of age. (B) Same as (A) for RPL13A-GFP. (C) Normalized growth rate for 4 classes of age. (D) Same as (A) for RPA190-GFP. (E) Same as (A) for SSF1-GFP. (F) From left to Right: pictures of a mother cell with ACT1pr-NLS-sfGFP, trapped in a cavity, before and after SEP. NLS-sfGFP total fluorescence and N/C ratio (calculated from NLS-sfGFP segmentation) as a function of generation, averaged after SEP alignment. Normalized N/C ratio for 4 classes of mother age. N= 14 cells. (G) Same as (F) for ACT1pr-sfGFP marker instead of ACT1pr-NLS-sfGFP. N= 28 cells. (H) From left to Right: pictures of a mother cell with VPH1-GFP, trapped in a cavity, before and after SEP. VPH1-GFP total fluorescence and vacuolar/cellular ratio as a function of generation after SEP alignment. Normalized vacuolar/cellular ratio for 4 classes of mother age. N= 26 cells. Scale bar: 5µm (for all pictures).

**Figure S3, related to Figure 5.**
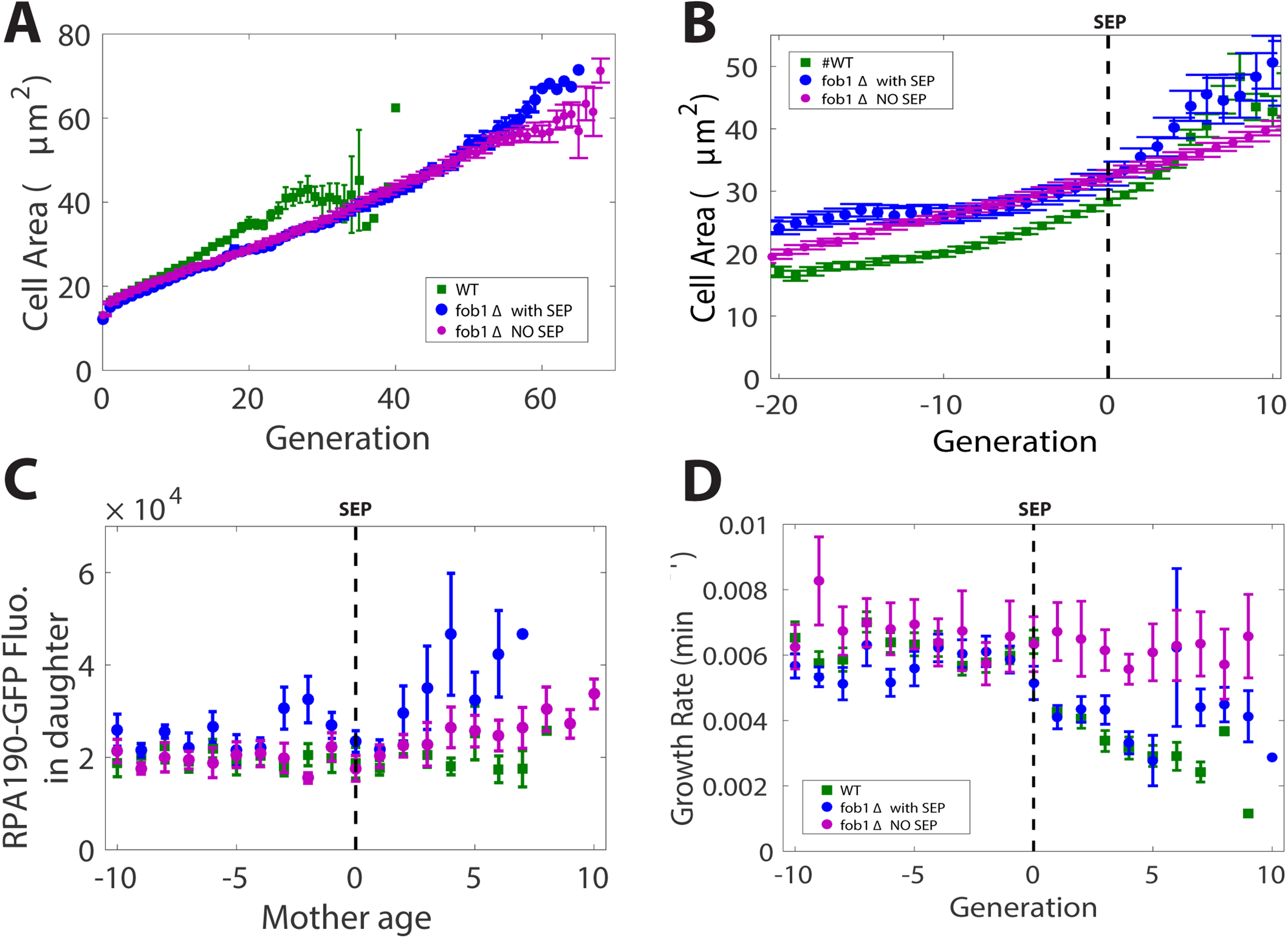
(A) Cell area in fob1Δ (N= 64 cells) and WT (N= 123 cells) after birth alignment. (B) Cell area in fob1Δ and WT after SEP alignment. (C) RPA190-GFP fluorescence in daughter cells for WT (green, N= 21 cells), “fob1 with SEP” (blue, N= 12 cells), “fob1 NO SEP” (magenta, N= 8 cells) after SEP alignment. (D) Growth rate (measured from projected area) as a function of generation, averaged after SEP alignment for WT (green, N= 36 cells), “fob1 with SEP” (blue, N= 22 cells), “fob1 NO SEP” (magenta, N= 18 cells).

**Movie S1, related to Figure1** Mother cell trapped in a cavity from birth to death and containing the fluorescence markers GFP-LacI (with LacO on each rDNA repeat) and Net1-mCherry. Time-lapse video. Left: phase contrast; middle: GFP; right: mCherry; cellular contour is delimited in yellow. Scale bar: 5µm.

**Movie S2, related to Figure2** Mother cell trapped in a cavity from birth to death and containing the fluorescence marker RPA190-GFP. Time-lapse video. Left: phase contrast; right: RPA190-GFP; cellular contour is delimited in yellow. Scale bar: 5µm.

**Movie S3, related to Figure2** Mother cell trapped in a cavity from birth to death and containing the fluorescence marker SSF1-GFP. Time-lapse video. Left: phase contrast; right: SSF1-GFP; cellular contour is delimited in yellow. Scale bar: 5µm.

**Movie S4, related to Figure3** Mother cell trapped in a cavity from birth to death and containing the fluorescence marker NOG2-GFP. Time-lapse video. Left: phase contrast; right: NOG2-GFP; cellular contour is delimited in yellow. Scale bar: 5µm.

**Movie S5, related to Figure3** Mother cell trapped in a cavity from birth to death and containing the fluorescence marker RPL13A-GFP. Time-lapse video. Left: phase contrast; right: RPL13A-GFP; cellular contour is delimited in yellow. Scale bar: 5µm.

**Movie S6, related to Figure3** Mother cell trapped in a cavity from birth to death and containing the fluorescence marker HTB2-sfGFP. Scale bar: 5µm.

**Movie S7, related to Figure5** fob1Δ mother cell, containing the fluorescence marker RPA190-GFP, which does not enter into senescence before death. Scale bar: 5µm.

**Movie S8, related to Figure5** fob1Δ mother cell, containing the fluorescence marker RPA190-GFP, which experiences SEP before death. Scale bar: 5µm.

## References

Brewer, B.J., and Fangman, W.L. (1988). A Replication Fork Barrier at the 3 ’End of Yeast Ribosomal RNA Genes. Cell 55, 637–643.

Defossez, P.-A., Prusty, R., Kaeberlein, M., Lin, S.-J., Ferrigno, P., Silver, P.A.,Keil, R.L.,and Guarente,L. (1999). Elimination of Replication Block Protein Fob1 Extends the Life Span of Yeast Mother Cells. Molecular Cell 3, 447–455.

Denoth-Lippuner, A., Krzyzanowski, M.K., Stober, C., and Barral, Y. (2014). Role of SAGA in the asymmetric segregation of DNA circles during yeast ageing. Elife 3.

Egilmez, N.K.,and Jazwinski, S.M. (1989). Evidence for the Involvement of a Cytoplasmic Factor in the Aging of the Yeast Saccharomyces cerevisiae. Journal of Bacteriology 171, 37–42.

Fehrmann, S., Paoletti, C., Goulev, Y., Ungureanu, A., Aguilaniu, H., and Charvin, G. (2013). Aging yeast cells undergo a sharp entry into senescence unrelated to the loss of mitochondrial membrane potential. Cell Rep 5, 1589–1599.

Gadal, O., Strauss, D., Petfalski, E., Gleizes, P.E., Gas, N., Tollervey, D., and Hurt, E. (2002). Rlp7p is associated with 60S preribosomes, restricted to the granular component of the nucleolus, and required for pre-rRNA processing. J Cell Biol 157, 941–951.

Ganley, A.R., Ide, S., Saka, K., and Kobayashi, T. (2009). The effect of replication initiation on gene amplification in the rDNA and its relationship to aging. Mol Cell 35, 683–693.

Ganley, A.R.,and Kobayashi, T. (2014). Ribosomal DNA and cellular senescence: new evidence supporting the connection between rDNA and aging. FEMS Yeast Res 14, 49–59.

Goulev, Y., Morlot, S., Matifas, A., Huang, B., Molin, M., Toledano, M.B.,and Charvin, G. (2017). Nonlinear feedback drives homeostatic plasticity in H2O2 stress response. Elife 6.

Ide, S., Saka, K., and Kobayashi, T. (2013). Rtt109 prevents hyper-amplification of ribosomal RNA genes through histone modification in budding yeast. PLoS Genet 9, e1003410.

Janssens, G.E., Meinema, A.C., Gonzalez, J., Wolters, J.C., Schmidt, A., Guryev, V., Bischoff, R., Wit, E.C., Veenhoff, L.M., and Heinemann, M. (2015). Protein biogenesis machinery is a driver of replicative aging in yeast. Elife 4, e08527.

Janssens, G.E., and Veenhoff, L.M. (2016). The Natural Variation in Lifespans of Single Yeast Cells Is Related to Variation in Cell Size, Ribosomal Protein, and Division Time. PLoS One 11, e0167394.

Jorgensen, P., Edgington, N.P., Schneider, B.L., Rupes, I., Tyers, M., and Futcher, B. (2007). The Size of the Nucleus Increases as Yeast Cells Grow Mol Biol Cell 18, 3523–3532.

Kennedy, B.K., Austriaco Jr., N.R., and Guarente, L. (1994). Daughter Cells of Saccharomyces cerevisiae from Old Mothers Display a Reduced Life Span The Journal of Cell Biology 127, 1985–1993.

Kobayashi, T. (2003). The Replication Fork Barrier Site Forms a Unique Structure with Fob1p and Inhibits the Replication Fork. Molecular and Cellular Biology 23, 9178–9188.

Kobayashi, T. (2008). A new role of the rDNA and nucleolus in the nucleus—rDNA instability maintains genome integrity. BioEssays 30, 267–272.

Kobayashi, T., Heck, D.J., Nomura, M., and Horiuchi, T. (1998). Expansion and contraction of ribosomal DNA repeats in Saccharomyces cerevisiae: requirement of replication fork blocking (Fob1) protein and the role of RNA polymerase I. Genes & Dev 12, 3821–3830.

Kressler, D., Hurt, E., and Bassler, J. (2017). A Puzzle of Life: Crafting Ribosomal Subunits. Trends Biochem Sci 42, 640–654.

Kume, K., Cantwell, H., Neumann, F.R., Jones, A.W., Snijders, A.P., and Nurse, P. (2017). A systematic genomic screen implicates nucleocytoplasmic transport and membrane growth in nuclear size control. PLoS Genet 13, e1006767.

Lu, K.L., Nelson, J.O., Watase, G.J., Warsinger-Pepe, N., and Yamashita, Y.M. (2018). Transgenerational dynamics of rDNA copy number in Drosophila male germline stem cells. Elife 7.

Miyazaki, T., and Kobayashi, T. (2011). Visualization of the dynamic behavior of ribosomal RNA gene repeats in living yeast cells. Genes Cells 16, 491–502.

Mortimer, R.K., and Johnston, J.R. (1959). Life Span of Individual Yeast Cells. Nature 183, 1751–1752.

Nakaoka, H., and Wakamoto, Y. (2017). Aging, mortality, and the fast growth trade-off of Schizosaccharomyces pombe. PLoS Biol 15.

Neumann, F.R., and Nurse, P. (2007). Nuclear size control in fission yeast. J Cell Biol 179, 593–6001.

Nishimura, K., Kumazawa, T., Kuroda, T., Katagiri, N., Tsuchiya, M., Goto, N., Furumai, R., Murayama, A., Yanagisawa, J., and Kimura, K. (2015). Perturbation of ribosome biogenesis drives cells into senescence through 5S RNP-mediated p53 activation. Cell Rep 10, 1310–1323.

Saka, K., Ide, S., Ganley, A.R., and Kobayashi, T. (2013). Cellular senescence in yeast is regulated by rDNA noncoding transcription. Curr Biol 23, 1794–1798.

Shcheprova, Z., Baldi, S., Frei, S.B., Gonnet, G., and Barral, Y. (2008). A mechanism for asymmetric segregation of age during yeast budding. Nature 454, 728–734.

Sinclair, D.A., and Guarente, L. (1997). Extrachromosomal rDNA Circles— A Cause of Aging in Yeast. Cell 91, 1033–1042.

Spivey, E.C., Jones, S.K., Rybarski, J.R., Saifuddin, F.A., and Finkelstein, I.J. (2017). An aging-independent replicative lifespan in a symmetrically dividing eukaryote. Elife 6.

Takeuchi, Y., Horiuchi, T., and Kobayashi, T. (2003). Transcription-dependent recombination and the role of fork collision in yeast rDNA. Genes Dev 17, 1497–1506.

Tiku, V., Jain, C., Raz, Y., Nakamura, S., Heestand, B., Liu, W., Spath, M., Suchiman, H.E.D., Muller, R.U., Slagboom, P.E.,et al. (2016). Small nucleoli are a cellular hallmark of longevity. Nat Commun 8, 16083.

